# Molecular Basis of Core Fucosylation-Dependent Modulation of IgG1 Fc–CD16a Binding

**DOI:** 10.64898/2026.06.26.732000

**Authors:** Natesan Mani, Alla Polozova, Srirupa Chakraborty

**Affiliations:** Department of Chemical Engineering, Northeastern University, Boston, MA 02115; Pivotal Attribute Sciences, Process Development, Amgen, Cambridge, MA 02141; Department of Chemistry and Chemical Biology, Northeastern University, Boston, MA 02115; Department of Physics, Northeastern University, Boston, MA 02115

**Keywords:** IgG1 Fc glycoengineering, Core fucosylation, FcγRIIIa/CD16a recognition, Molecular dynamics simulations, Glycan-mediated conformational dynamics

## Abstract

Core fucosylation of the IgG1 Fc N297 glycan is known to reduce binding affinity to the FcγRIIIa (CD16a) receptor and attenuate antibody-dependent cellular cytotoxicity (ADCC), yet the structural mechanisms underlying this effect remain incompletely understood. Here, we use extensive all-atom molecular dynamics simulations to systematically investigate how Fc glycosylation modulates the structural, energetic, and dynamical landscape of the IgG1 Fc–CD16a complex across multiple systems with fucosylation and galactosylation. Relative binding free energy calculations reproduce experimentally established trends, showing that afucosylation consistently strengthens Fc–CD16a interactions. Mechanistically, dual fucosylation (on both Fc arms) increases inter-glycan packing between the Fc N297 glycans, restricts Fc glycan conformational sampling, and destabilizes the conformational organization of the CD16a N162 glycan. These glycan-mediated perturbations propagate to the protein interface. The result is reduced Fc–CD16a contact persistence, redistribution of energetically important residues away from the canonical binding interface, and broader, less stable receptor-bound conformational states. Dynamic cross-correlation analysis further reveals that afucosylated systems maintain substantially stronger coordinated motions across the Fc–CD16a assembly, whereas fucosylation disrupts long-range dynamic coupling between the receptor and antibody domains. Across these different energetic, structural, conformational, and dynamical readouts, fucosylation systematically shifts the Fc–CD16a assembly from a compact, interface-stabilized binding mode toward a more heterogeneous and weakly coupled receptor-bound ensemble. Together, our findings set forth a mechanistic basis for Fc glycosylation regulating receptor engagement through ensemble-level conformational and dynamical reorganization rather than simple local steric effects. These results provide mechanistic design principles for rational Fc glycoengineering and the development of therapeutic antibodies with enhanced effector functions. More broadly, this work highlights how glycan composition can be leveraged as a tunable molecular design parameter for engineering protein recognition, conformational stability, and immune effector function in therapeutic glycoproteins.

## INTRODUCTION

Adaptive immunity, which is acquired from antibody-based therapies, plays a key role in defending the body from pathogenic agents and in regulating antibody production^1^. The IgG (Immunoglobulin G) is the most prevalent antibody, and the IgG1 antibody is its most prevalent subtype. IgG possesses two domains: the Fragment antigen binding domain (Fab), responsible for antigen recognition and the Fragment crystallizable (Fc) domain, which interacts with immune cells to induce potent effector functions^2^. The Fc region of the IgG1 is in charge of eliciting effector function from antibodies by binding to the FcγR receptor present on the surface of immune cells^3^. This, in turn, is responsible for phagocytosis, antibody-dependent cellular cytotoxicity (ADCC), and regulation of antibody production^4^. FcγRIII (or CD16), a subclass of FcγR, binds to the IgG1 Fc with a strong affinity^5^, which is further influenced by the glycosylation profile on both the Fc of IgG1 and the FcγRIII glycoproteins^5, 6^. FcγRIIIa (CD16a), a subtype of FcγRIII, is predominantly present on the surface of natural killer (NK) cells and monocytes and hence is a primary target to elicit an immune response (see **Figure 1A** for a schematic overview). Fc glycosylation is a key determinant of IgG1 antibody function, yet the molecular mechanisms by which specific glycan features modulate receptor binding remain incompletely understood. In particular, while core fucosylation is known to dramatically reduce Fc–CD16a affinity, the structural and dynamic origins of this effect are still not well elucidated. Understanding how specific glycan motifs regulate receptor binding through conformational and dynamical control is an important challenge in the molecular engineering of therapeutic glycoproteins.

**Figure 1:**
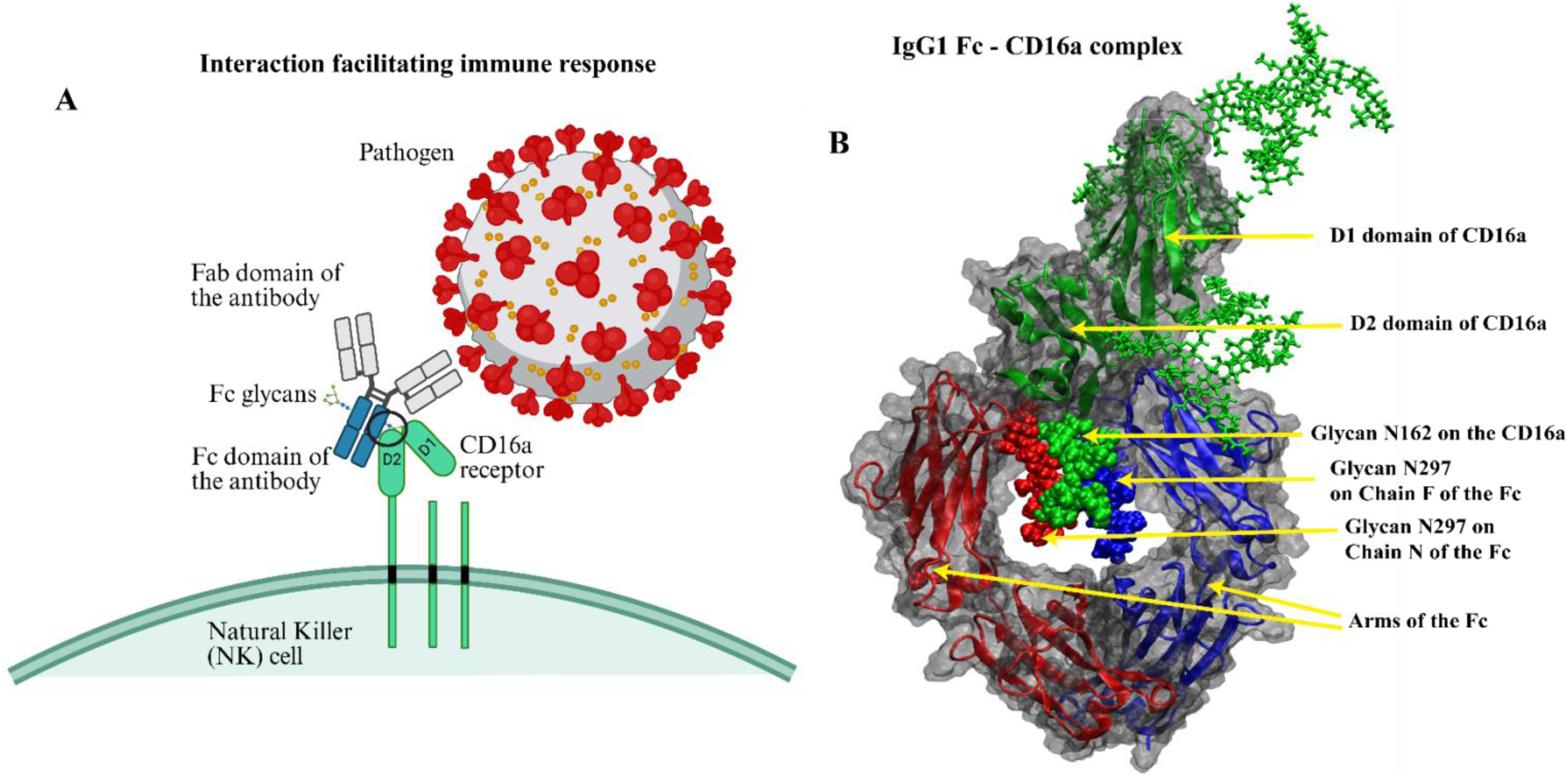
A pictorial overview of the innate immune response and the structure of the IgG1 Fc - CD16a complex with modeled N-linked glycans. The interactions facilitating the immune response are shown on **Panel A**. The Fab domain of the antibody is involved in antigen/pathogen recognition and binding while the Fc domain interacts with receptors such as CD16a on the surface of NK cells and phagocytes to facilitate immune response. The glycans are also shown on the Fc, which are known to play a key role in determining the strength of binding and hence the extent of immune response. The overall structure of the bound, simulated domains of the IgG1 Fc - CD16a complex (see circled region in **Panel A**), is shown in **Panel B**. The Fc region consists of two glycans present at residue N297 and the CD16a domain is shown in green which has two sub-domains D1 and D2, which are glycosylated based on the most probable N-linked glycans obtained from mass spectroscopy data (see Figure 2 for more information).

The Fc region of the IgG1 has two N glycosylation sites, one on each arm, located at residue N297, which are highly conserved. The glycans modulate the structure and conformation of the Fc, which helps in its binding to the FcγR receptor^3, 7^. The extracellular domain of CD16a contains two domains, D1 and D2, held together by a flexible linker. The D1 domain contains the primary glycosylation sites, and these glycans can influence the receptor’s stability, expression, and interaction with Fc. The D2 domain is directly involved in the binding to the Fc, and its structure allows for the proper alignment and binding^8^. There are a total of five glycans on the receptor, at positions N38, N45, N74, N162, and N169^9^. Glycans N38, N45, and N74 are on the D1 domain and, as mentioned above, influence the receptor’s stability. Glycans N162 and N169 are present on the D2 domain, and N162 plays a pivotal role in the binding to the Fc of IgG1^10, 11^. Although all receptor glycans were modeled to account for their potential long-range effects on structure and interaction energetics, N162 and N169 are of special interest because of their proximity to the Fc-binding interface and likely their more direct role in modulating local contacts. **Panel B** of **Figure 1** shows the overall structure of the simulated Fc–CD16a complex, with the two N297 glycans on the Fc and the D1 and D2 sub-domains of CD16a labelled.

A key strategy for engineering stronger interactions between IgG1 and the CD16a receptor is the rational control of Fc glycosylation, where subtle changes in glycan composition can substantially alter antibody binding, conformational stability, and downstream immune effector responses. Among these modifications, the most well-established is the removal of core fucose from the Fc N-glycan, which substantially increases CD16a binding affinity^1, 10, 12–14^. This stronger Fc–CD16a interaction enhances antibody-dependent cellular cytotoxicity (ADCC), allowing effector responses to be triggered at lower IgG1 concentrations and highlighting Fc glycoengineering as a practical route to improve therapeutic antibody potency^13, 15^. There is some evidence that suggests that the N162 glycan on CD16a forms stabilizing contacts with the Fc N297 glycan, and that core fucose disrupts these interactions^15^. Removal of fucose is therefore thought to relieve local steric interference and promote more favorable Fc–CD16a engagement. However, the molecular origin of this effect is not fully resolved. Beyond direct steric effects, afucosylation is likely to alter glycan conformational sampling and longer-range interfacial coupling, which are difficult to dissect experimentally because of glycan heterogeneity and dynamics. This raises the possibility that Fc glycosylation acts not merely as a local structural modification, but as a tunable regulator of the conformational ensemble and dynamic communication network governing receptor engagement. Other Fc glycan features, including terminal galactosylation, have also been reported to enhance CD16a binding^1, 13, 16^, and the effects of galactose and absence of fucose may be additive^17, 18^ Nevertheless, the impact of galactosylation is generally smaller than that of afucosylation, making core fucose removal the most direct, robust, and experimentally practical strategy for engineering high-affinity Fc variants.

Despite clear experimental evidence that core fucosylation of the IgG1 Fc N297 glycan strongly reduces CD16a binding affinity and downstream effector function, the molecular origins of this effect remain difficult to resolve because glycans are structurally heterogeneous, highly flexible, and capable of modulating recognition through both local contacts and longer-range dynamic coupling. In particular, it remains unclear whether the detrimental effects of fucose arise primarily from direct steric interference at the Fc–CD16a interface or from broader conformational and dynamical reorganization of the receptor-bound complex. These challenges motivate a simulation-based framework capable of directly interrogating how defined fucosylation states reshape the energetic and conformational landscape of the IgG1 Fc–CD16a assembly.

Accordingly, in this study, we systematically investigate the structural and dynamical consequences of Fc core fucosylation across a panel of Fc glycoforms with varying fucosylation and galactosylation states. The IgG1 antibody was selected because of its prevalence in human serum and central importance in therapeutic antibody engineering, while CD16a was chosen due to its strong role in mediating Fc-dependent immune effector responses. Using extensive all-atom molecular dynamics simulations consisting of three independent ∼1.2 μs trajectories per system, we characterize how core fucosylation alters binding energetics, glycan organization, protein-protein contacts, conformational landscapes, and long-range dynamic coupling across the Fc–CD16a interface. These results provide a mechanistic framework for understanding how core fucosylation regulates Fc receptor recognition and establish broader design principles for rational Fc glycoengineering and therapeutic antibody optimization. Resolving these molecular effects provides a pathway toward engineering glycan-controlled conformational ensembles and tuning antibody effector functions through precise control of Fc glycosylation. More broadly, this work helps to establish how glycan composition can be leveraged as a tunable molecular design parameter for engineering biomolecular recognition and immune signaling.

## RESULTS

### Glycoform Design and Simulation Overview

The MD simulations were performed on a panel of 6 IgG1 Fc glycoforms designed to systematically probe the effects of Fc N-glycan composition. These glycoforms consist of paired biantennary afucosylated (expected higher affinity) and paired biantennary fucosylated (expected lower affinity) systems. From literature, we also know that the CD16a receptor binds asymmetrically to the two chains of the Fc^1, 19^. This is reflected in our study by assuming Chain N to be the arm of the Fc closer to the receptor (or the Near chain), and Chain F to be the distal arm (or the Far chain). **Figure 2** and **SI Table 1** illustrate the different types of glycans, and systems modelled on the Fc, and **Figure 2** also represents a structural view of each individual glycosylated domain, along with the bound structure. The glycosylation on the CD16a was modeled based on the most probable glycans on each site, based on mass spectroscopy data^20^, with the N162 glycan represented by High mannose (M5) to elicit the strongest binding^9, 12^. The crystal structure of the Fc–CD16a pair (code: 1E4K) was obtained from RCSB^19^. Additionally, different galactosylation levels on the glycan are represented by G0 (no galactose residues), G1 (single galactose residue) and G2 (two galactose residues). The final list of systems is represented in **SI Table 1**. Three independent runs were then performed on AMBER using the CHARMM36m forcefield^21^, each for 1.2µs. For the analysis performed, the first 200 ns of all the trajectories were discarded to ensure that the systems had converged. Details of system setup and simulation protocol can be found in the **Methods** section.

**Figure 2:**
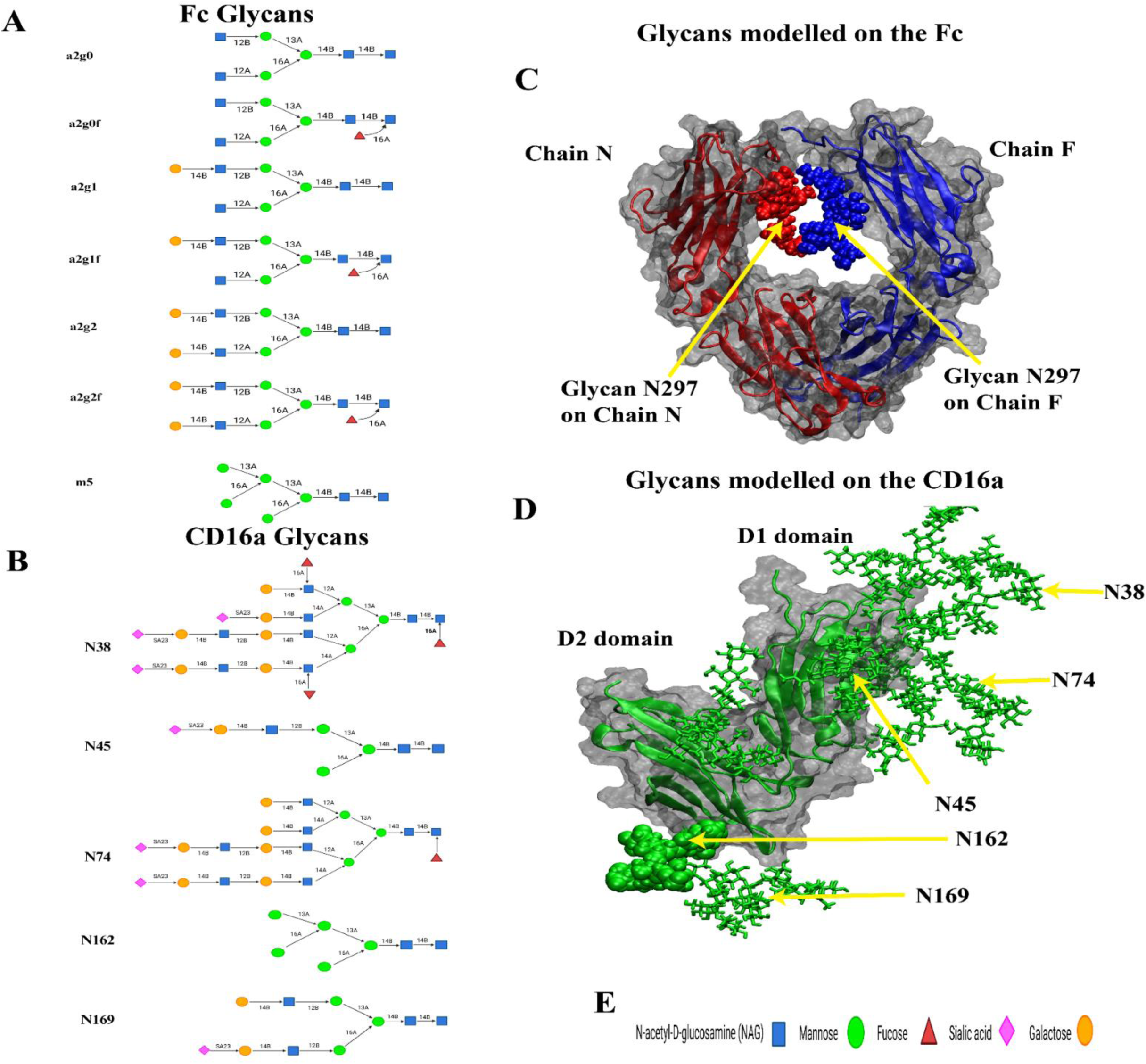
Representation of the glycans modeled on the Fc and the CD16a. In the above figure, **Panels A** and **C** shows he different types of glycans that were used to model the N297 glycans on the Fc, based on most commonly observed tructures in IgG antibodies and different degrees of fucosylation. The standardized Symbolic Nomenclature for Glycans SNFG) representation (see **Panel E**) has been used, **Panels B** and **D** show the glycans modelled on the possible sites of he CD16a based on published mass spectroscopy data from Jacob et al.

### Afucosylation enhances Fc–CD16a interactions in agreement with experiments

Experimental studies have consistently shown that removal of the core fucose from the IgG1 Fc N297 glycan enhances binding to CD16a, thus increasing antibody-dependent cellular cytotoxicity^1, 10, 12–14, 22^. This enhancement has been proposed to arise from improved glycan-mediated stabilization and more favorable intermolecular packing at the Fc–CD16a interface, particularly involving interactions between the Fc glycans and the N162 glycan of CD16a^15^. Hence, removing fucose from the N297 glycan is a proven strategy to stabilize the Fc–CD16a complex, leading to stronger binding. A key objective of this study was therefore to determine whether our simulation framework could reproduce these experimentally established directional changes in binding energetics across different Fc glycoforms.

To assess relative interaction strengths across the simulated systems, we computed a Molecular Mechanics / Poisson-Boltzmann Surface Area (MM/PBSA) style interaction energy estimate from the MD trajectories using AmberTools-based energetic decomposition^23, 24^. The Van der Waals and Electrostatic interaction terms were empirically scaled (α = 0.158 and β = 0.153, respectively, see **Methods**), following the approach of Minghui et al^25^. This approach was to obtain results that are on a comparable scale to experimental readouts. Importantly, these calculations were not intended to provide absolute binding free energies, but rather to quantify comparative energetic changes associated with glycan composition. Our simulations successfully reproduced the expected enhancement in Fc–CD16a interaction strength when the Fc N297 glycans are afucosylated (see **Figure 3**). The agreement of these directional energetic trends with experimental observations provides an important validation of the underlying conformational ensembles and intermolecular interaction networks sampled in the simulations. This validation is particularly significant because it suggests that the simulations capture the relevant molecular determinants governing glycan-dependent modulation of Fc–CD16a recognition.

**Figure 3:**
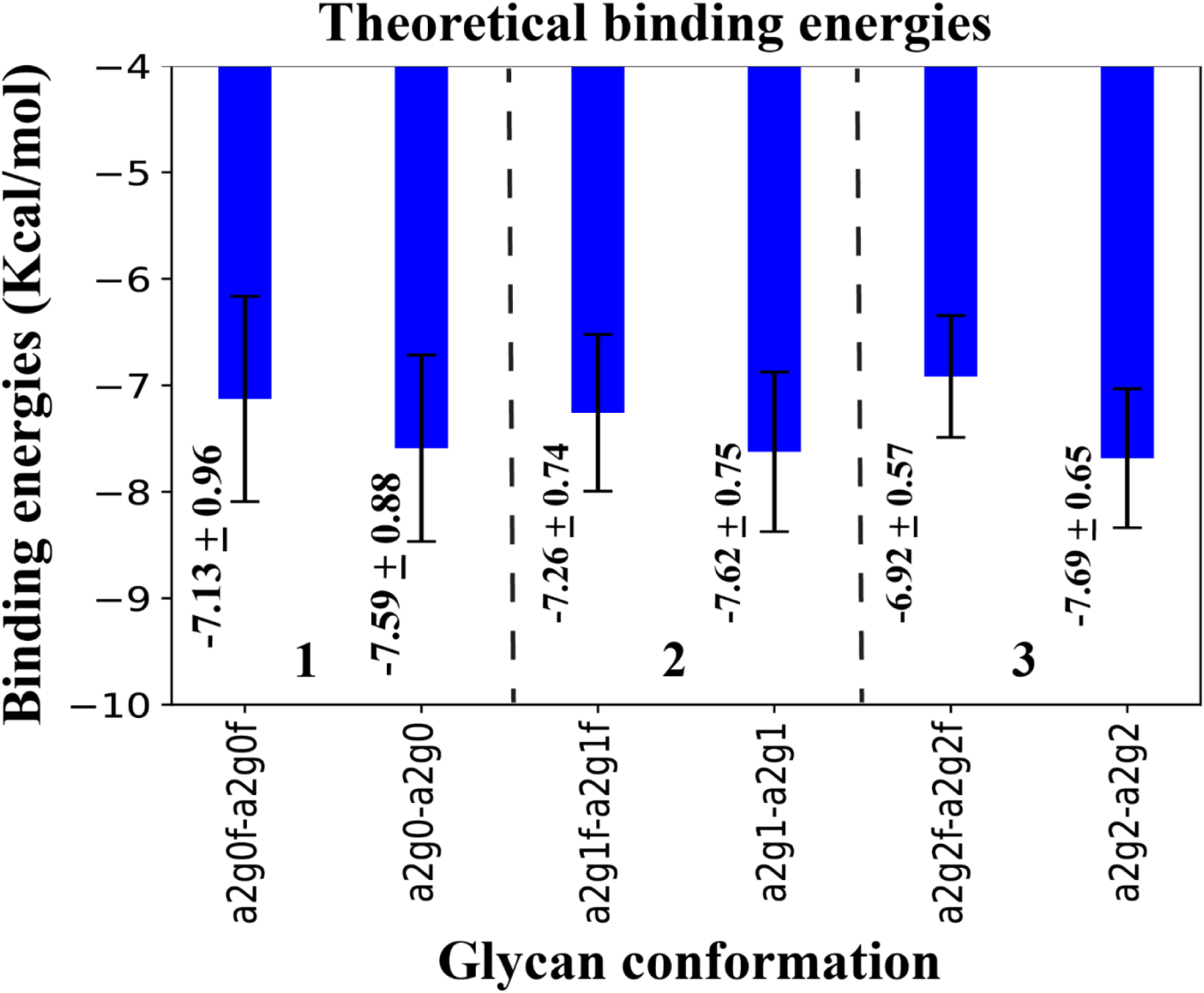
Removal of fucose increases binding affinity. Binding energy analysis using empirically scaled MMPBSA, comparing fucosylated (left) and afucosylated (right) systems for each glycan pair (enclosed between dotted lines). Binding energy increased markedly upon fucose removal, as seen in **Blocks 1, 2** and **3.**

### Fucosylation modulates Fc glycan packing and CD16a N162 dynamics

Having established that the simulations reproduce experimentally observed afucosylation-dependent binding trends, we next investigated the structural mechanisms underlying these energetic differences. Literature suggests that the formation of the Fc–CD16a complex is not only driven by protein-protein interactions but is also mediated by glycan-protein and glycan-glycan interactions^26–29^. To understand these glycan contributions, we examined interactions between the two Fc N297 glycans for the paired biantennary afucosylated and paired biantennary fucosylated glycoforms by quantifying heavy-atom contacts between the glycans across the trajectories. Fucosylation resulted in a net increase in total inter-glycan contacts (**Figure 4A-B**). An exception was observed for G2 glycoforms, where the afucosylated case exhibited higher contact counts (**Figure 4C**), likely owing to the added stability conferred by the terminal galactose residues. Nevertheless, the overall trend was consistent; inter-glycan distances remained shorter in the fucosylated systems. These increased contacts suggest that the presence of core fucose promotes tighter Fc glycan packing and restricts glycan conformational sampling. Such constrained glycan organization may reduce the ability of the Fc glycans to adopt receptor-oriented conformations that stabilize the Fc–CD16a interface. Our next step was to look at possible contacts that are being formed between the Fc glycans and the glycan N162 on the CD16a, which has been previously implicated in Fc-receptor recognition^11, 12, 15, 16, 29–31^. Surprisingly, we did not observe persistent direct glycan–glycan contacts across the trajectories. Our observation suggests that the effects of fucosylation may instead arise indirectly through altered glycoprotein conformational dynamics, as has been indicated in some prior studies^15, 16, 20, 26, 28, 29, 32^. In light of this, we looked at the conformational motion of the N162 glycans by plotting Free Energy Surface (FES) plots (see **Methods**). We used two reaction coordinates to capture these conformations: Radius of gyration (R_g_) and the distance between the Centre of Masses (COM) of the N162 glycan and the Fc glycans (**Figure 4D-F**). We observe that in paired biantennary fucosylated systems, the N162 glycan samples a substantially broader conformational space, indicating that the presence of dual fucosylation disrupts the interactions, leading to a reduction in the stabilizing contacts that it makes with both the Fc glycans and the Fc protein. Together, these results indicate that fucosylation restricts Fc glycan dynamics while simultaneously destabilizing the N162 glycan conformation, ultimately contributing to the reduced binding affinity observed in **Figure 3**.

**Figure 4:**
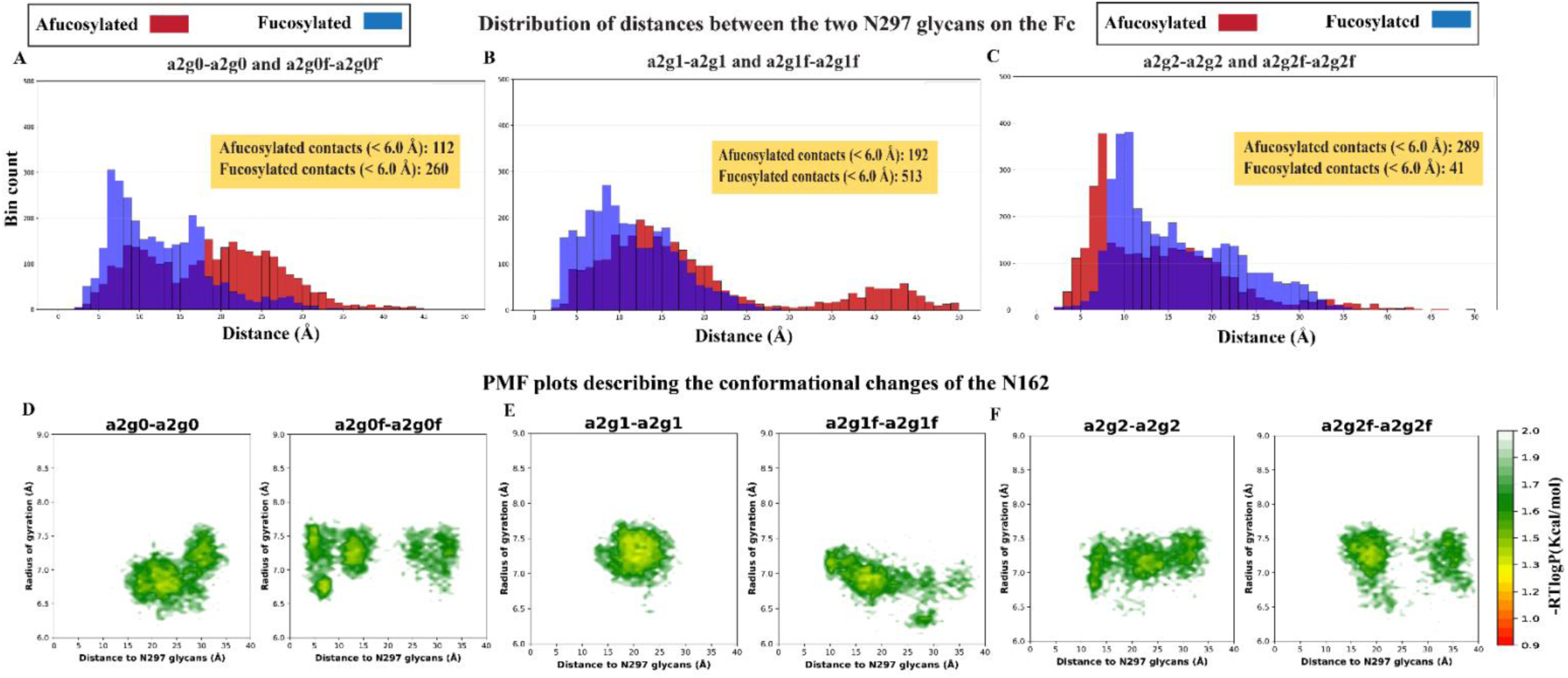
Fucosylation increases Fc glycan contacts, while destabilizing the CD16a N162-glycan conformation. Inter-glycan contact analysis between the two Fc N297 glycans (**Panels A–C**) reveals substantially higher contact frequencies in Paired biantennary fucosylated systems (blue) compared to the Paired biantennary afucosylated systems (red). This is observed for G0 (**Panel A**) and G1 (**Panel B**) systems. For G2 (**Panel C**), the afucosylated system shows a higher number of contacts, potentially due to added stability from terminal galactose residues; however, the overall inter-glycan distances remain shorter in the fucosylated systems relative to the afucosylated systems. These increased inter-glycan contacts in the fucosylated systems likely promote tighter Fc glycan packing and restrict N297 glycan conformational sampling, whereas afucosylated glycans access a broader conformational ensemble. Mean ± SD inter-glycan distances for each panel are: G0 (afucosylated = 18.69Å ± 2.48Å, fucosylated = 12.48Å ± 0.62Å), G1 (afucosylated = 18.80Å ± 4.43Å, fucosylated = 11.19Å ± 1.86Å) and G2 (afucosylated = 13.91Å ± 4.94Å, fucosylated = 16.11Å ± 3.97 Å). Conversely, fucosylation destabilizes the N162 glycan of the CD16a receptor, as shown by the Free Energy Surface (FES) plots representing two reaction coordinates: (i) distance between the N162 glycan and the N297 glycans on the Fc, and (ii) radius of gyration (Rg) of the N162 glycan. We observe an expanded conformational space sampled by the fucosylated glycans relative to their afucosylated counterparts for G0 (**Panel D**), G1 (**Panel E**) and G2 (**Panel F**). Together, these divergent effects of fucosylation, compacting Fc glycans while destabilizing the CD16a N162-glycan, reduce productive glycan-glycan contacts and compromise stable Fc–CD16a complex formation.

### Afucosylation stabilizes the Fc–CD16a protein-protein interaction interface

Our previous analyses showed that core fucosylation alters Fc glycan organization and increases conformational heterogeneity of the CD16a N162 glycan. We next investigated whether these glycan-mediated effects propagate to the protein interface itself by examining protein–protein contact frequencies between the IgG1 Fc and CD16a receptor. To quantify differences in interfacial stability, residue-level protein–protein contact frequencies were computed across the trajectories using a distance-based contact definition (see **Methods**). To highlight the effect of fucosylation, differential contact maps were generated, and the contact frequency difference values were calculated by subtracting the fucosylated values from the afucosylated values. Positive values, therefore, indicate contacts that are more frequent or persistent in the afucosylated complexes. We find that paired biantennary afucosylated systems exhibited substantially stronger and more persistent Fc–CD16a contacts relative to their fucosylated counterparts, as evidenced by the widespread enrichment of positive contact frequency differences (see **Figure 5A-C**). These observations indicate that core fucosylation destabilizes optimal interfacial packing between the Fc and CD16a receptor. This observation is consistent with our earlier findings (see **Figure 4**) that dual fucosylation strongly perturbs Fc glycan organization and N162 conformational dynamics. Together, these results support a coupled mechanism in which fucosylation-induced glycan rearrangements indirectly weaken the Fc–CD16a protein interaction profile, leading to less stable receptor engagement.

**Figure 5:**
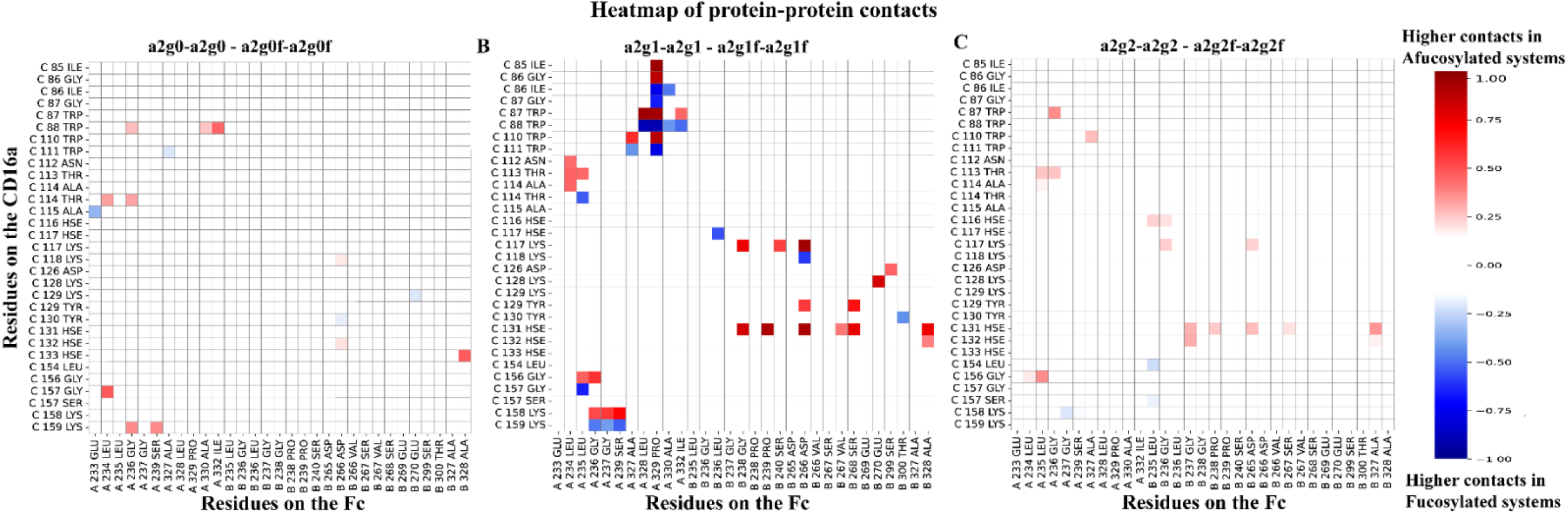
Afucosylated systems form more persistent protein-protein contacts than fucosylated systems. Contact frequency analysis reveals substantial differences between Paired biantennary afucosylated and fucosylated systems. The fucosylated complex exhibits significantly reduced contact frequencies compared to its afucosylated counterpart. This can be seen across varying degrees of galactosylation, ranging from G0 (see **Panel A**), G1 (see **Panel B**), and G2 (see **Panel C**), indicating that fucose residues hinder productive protein-protein interactions. These results demonstrate that fucosylation allosterically modulates the Fc-CD16a interface by directly suppressing critical contacts necessary for stable complex formation.

### Fucosylation redistributes interfacial energetics and leads to a sub-optimal orientation of the CD16a

Having established that core fucosylation weakens Fc–CD16a contacts and perturbs glycan conformational organization, we next investigated how these effects influence the energetic organization of the binding interface. To characterize residue-level energetic contributions, we performed per-residue decomposition of the MM/PBSA-derived interaction energies and identified the dominant residues contributing to Fc–CD16a stabilization in each glycoform system. The top 10 residues on both the Fc and the CD16a that contribute to observed energies are selected. To emphasize the energetic differences between fucosylated and afucosylated systems, only the residues that are uniquely enriched in these species are mapped onto the structure (see **Figure 6 and SI Table 2**). Across the different paired biantennary glycoforms, we find that the afucosylated systems displayed energetically dominant residues concentrated near the canonical Fc–CD16a binding interface, on the D2 domain of the receptor (**Figure 6A–C**), consistent with a more favorable and tightly organized interfacial arrangement. In contrast, the paired biantennary fucosylated systems exhibited a marked redistribution of energetically significant residues toward the D1 domain of CD16a, rather than the interface-proximal D2 domain (see circled regions in **Figure 6D-F**). The emergence of these energetically active residues “farther” away from the binding interface indicates a conformational shift induced by the presence of the two fucose residues on the Fc, consistent with a less optimal interfacial arrangement. These findings, along with the previous observation of reduced protein-protein contacts, suggest that core fucosylation does not simply weaken isolated interfacial contacts but instead reorganizes the energetic landscape of the Fc–CD16a complex, leading to altered receptor positioning and reduced binding stability.

**Figure 6:**
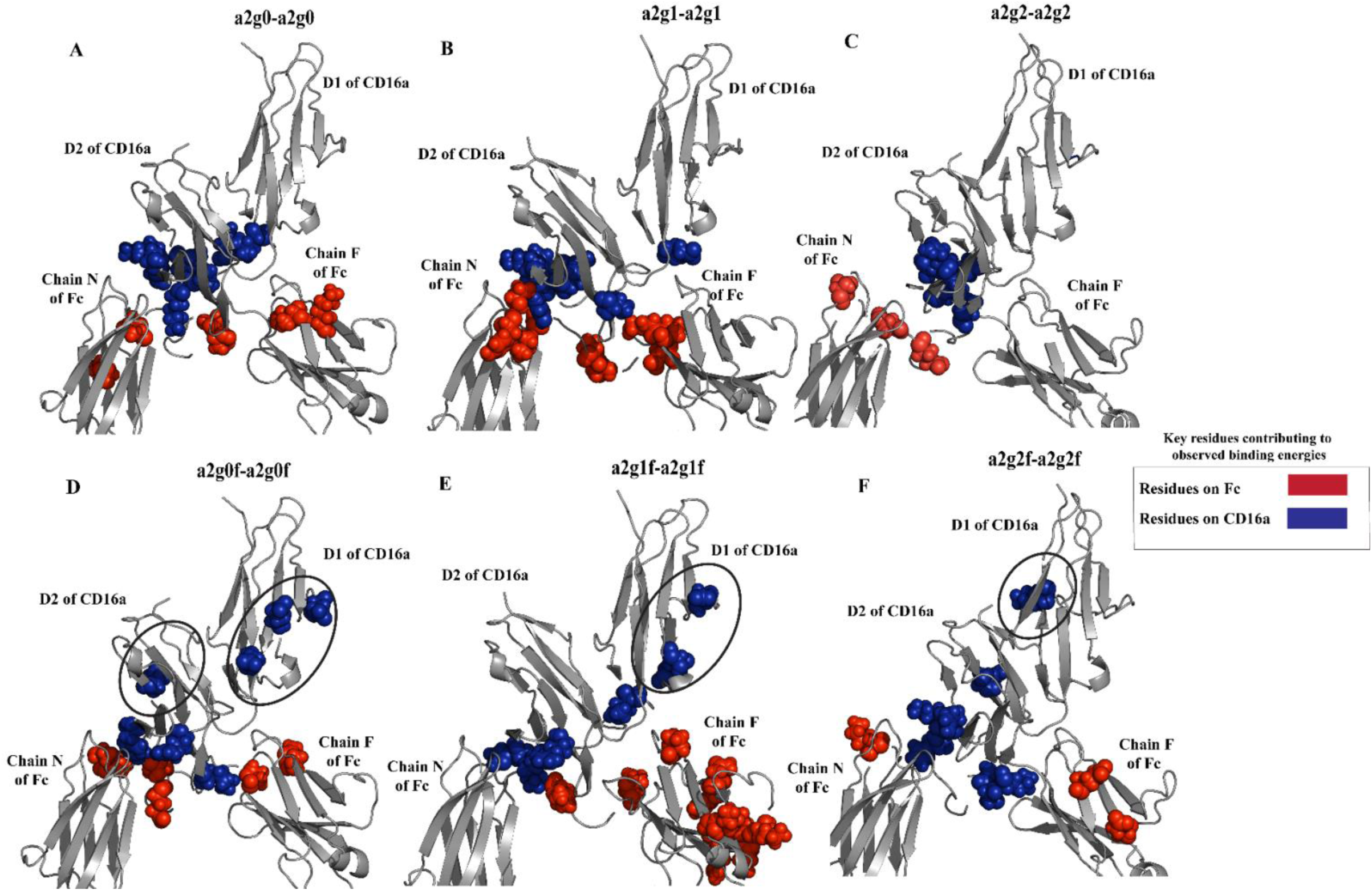
Fucosylation alters the energetic landscape of the binding complex. Energy decomposition analysis identified the 20 most energetically active residues at the Fc-CD16a interface that are unique to Paired biantennary ucosylated and afucosylated systems. This was done for varying degrees of galactose addition, ranging from G0 (**Panels A** and **D**), G1 (**Panels B** and **E**), and G2 (**Panels C** and **F**) and the list of residues can be seen in **SI Table 2**. We find that energetically critical residues on CD16a in the fucosylated complex (circled) are spatially displaced from the binding interface compared to the afucosylated system, indicating that fucosylation fundamentally alters the energetic landscape of the interaction.

### Fucosylation destabilizes preferred Fc–CD16a conformational states

The observed weakening of protein-protein interactions and redistribution of interfacial energetics in the paired biantennary fucosylated systems prompted us to investigate whether core fucosylation also alters the global conformational landscape of the Fc–CD16a complex. To characterize receptor positioning relative to the Fc domain, we quantified CD16a rotational motion along two orthogonal axes of rotation, along with the distance between the Centre of masses (COM) of CH2 domains of the Fc (see **Figure**s **7G-H**). These collective coordinates were analyzed using two-dimensional Free Energy Surface (FES) landscapes, where deeper and more localized basins correspond to more stable and preferentially populated conformational states (see **Methods**).

**Figure 7:**
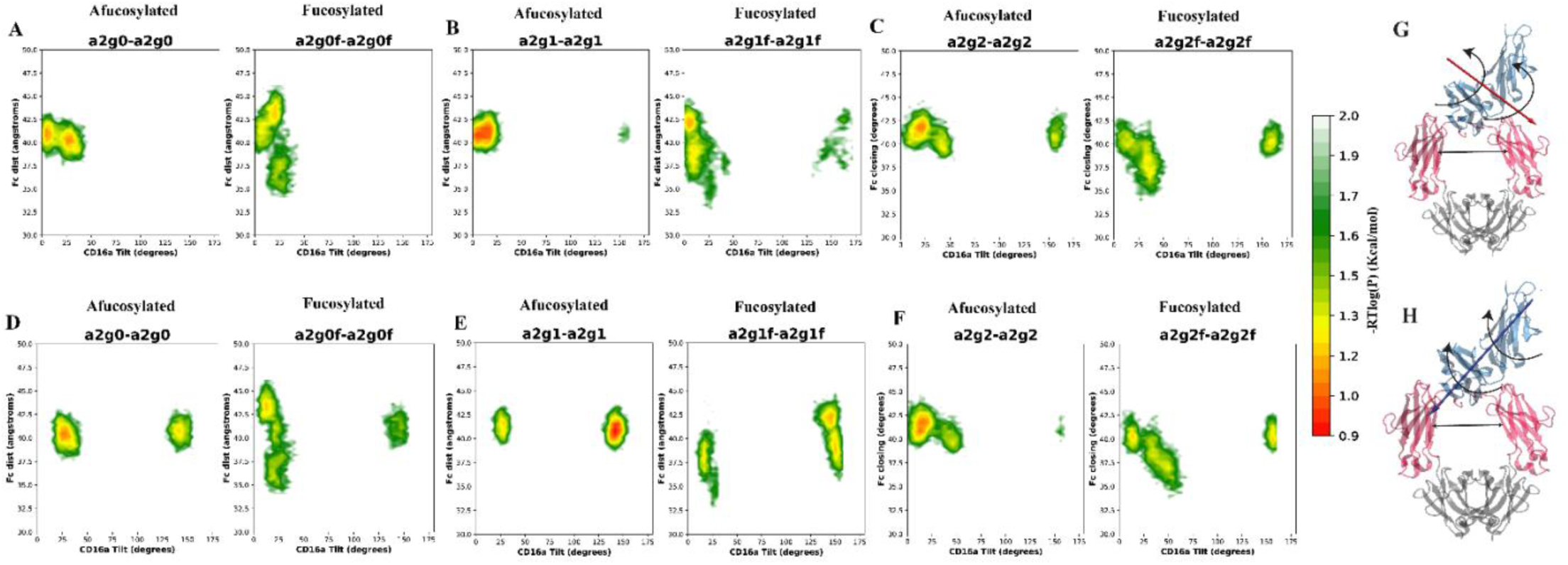
Fucosylation perturbs Fc–CD16a conformational stability. Free Energy Surface (**FES**) plots representing CD16a motion along two rotational axes (shown in **G** and **H**). Lower colorbar values indicate regions of higher stability. This analysis is performed for different galactosylation frequencies ranging from G0 (**Panels A** and **D**), G1 (**Panels B** and **E**) and G2 (**Panels C** and **F**). We observe that the Paired biantennary fucosylated systems sample a larger conformational space, while the afucosylated counterparts exhibit discrete regions of high stability characterized by compact, well-defined energy wells, and this is observed for all degrees of galactosylation and across both the rotational axes. This indicates that the CD16a receptor is locked in a binding accessible conformation in afucosylated species, while in fucosylated systems it is locked in a sub-optimal binding conformation, leading to a larger sampling of conformations.

Consistent with our previous findings, we found that the paired biantennary afucosylated species (see plots under **afucosylated** in **Figure 7**) exhibited compact and well-defined free-energy basins along both rotational coordinates, indicating preferential sampling of stable receptor-binding conformations. In contrast, the corresponding paired biantennary fucosylated systems sampled broader and more diffuse conformational landscapes, characterized by reduced basin localization and increased conformational heterogeneity (see plots under **Fucosylated** in **Figure 7**). The observed changes span both rotational motion of CD16a and variations in the Fc COM distance, suggesting coordinated alterations in receptor orientation and Fc domain geometry. Together, these results indicate that fucosylation shifts the conformational landscape of the Fc, which in turn alters CD16a positioning and promotes suboptimal binding configurations.

### Afucosylation enhances coordinated dynamics across the Fc–CD16a interface

Our previous analyses indicated that core fucosylation weakens Fc–CD16a binding, perturbs glycan organization, redistributes interfacial energetics, and destabilizes preferred receptor-binding conformations. Collectively, these findings suggested that the structural consequences of fucosylation propagate beyond local glycan rearrangements and influence larger-scale dynamic coordination across the complex. We therefore next investigated whether fucosylation alters correlated motions and dynamic coupling between the Fc and CD16a domains.

To characterize coordinated motions throughout the complex, we calculated the Dynamic Cross-Correlation Matrices (DCCM). DCCM is a method of quantifying time-correlated motions of each residue with all other residues throughout the protein^33^. We represented them as a two-dimensional matrix, and to have a clearer understanding of the impact brought about by fucose, we took the difference of the afucosylated and fucosylated matrices, respectively. On plotting these differences, we observed significantly stronger correlations in the afucosylated systems compared to the fucosylated systems (see **Figure 8**). These enhanced correlations were particularly prominent between the Fc domains themselves (black squares in **Figure 8**) and across the Fc–CD16a interface (red squares in **Figure 8**). Notably, the strongest increases in coordinated motion involved Fc chain N, which lies proximal to the receptor-binding region, and the D2 domain of CD16a, which directly participates in Fc recognition.

**Figure 8:**
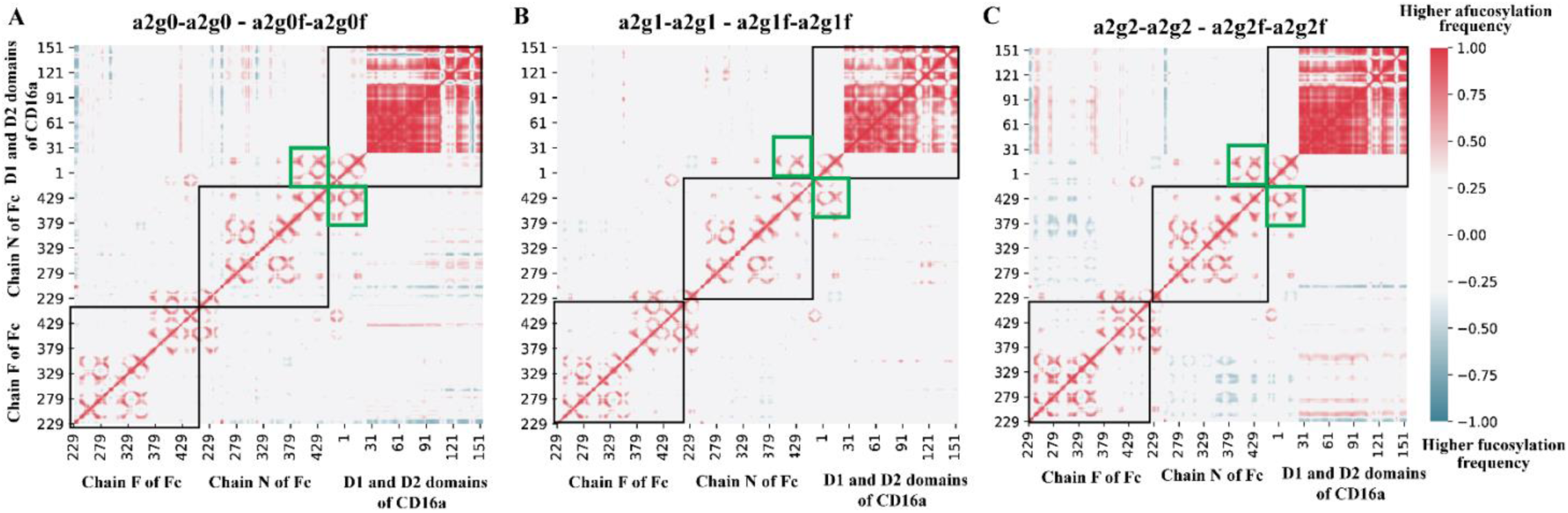
Core fucosylation disrupts dynamic coupling across the Fc–CD16a assembly. **Panels A, B,** and **C** represent three different pairs of glycans with different degrees of presence of galactose (G0, G1, and G2, respectively). The plots represent the difference in correlation frequencies between Paired biantennary afucosylated and fucosylated systems, with higher afucosylated frequencies shown in red and higher fucosylated frequencies in blue. Enhanced correlations in the afucosylated system are particularly evident in intra-Fc and intra-CD16a interactions (black squares) and across Fc-CD16a interface regions, most notably between Fc chain N (proximal to CD16a) and the CD16a D1 domain (green squares). These findings indicate that fucose removal enhances dynamic coupling within the Fc region and strengthens coordinated motions at the Fc-CD16a interface.

The enhanced dynamic coupling observed in the afucosylated systems suggests that removal of core fucose promotes more coordinated inter-domain motions across the Fc–CD16a complex. In contrast, fucosylation disrupts this dynamic coordination, leading to weaker coupling between the receptor and Fc domains. Together, these findings support a model in which core fucosylation weakens Fc receptor engagement not only through local structural perturbations, but also by disrupting collective motions that stabilize the receptor-bound state.

## DISCUSSION

In this study, we have investigated the role of fucosylation across a range of galactosylation levels and provided a structural and dynamical interpretation for the experimentally observed affinity differences. Together, these findings contribute toward a mechanistic framework for the rational glycoengineering of antibody therapeutics. Our simulations have revealed that across most of the glycosylation patterns, fucose causes a reduction in binding affinity, in agreement with literature findings (**Figure 3**). These findings led to two important questions: Are the observed binding affinity differences driven primarily by direct inter-glycan interactions, or do the glycans instead modulate the underlying protein-protein binding events? To address the first possibility, we examined the interactions between the two Fc glycans and found that fucosylation leads to a greater frequency of inter-glycan contacts, which indirectly constrains the motion of these two Fc glycans (**Figure 4A-C**). This increase in glycan–glycan contacts may, in part, arise from the conformational effects of core fucosylation itself. In our previous work, we showed that core fucosylation induces a characteristic bend in the orientation of N-glycans, altering the spatial disposition of the glycan branches^34^. Such a change in glycan orientation would naturally change the propensity for inter-glycan interactions between the Fc glycans. Concurrently, we also find that in the presence of dual fucose residues, the N162 glycan on the CD16a receptor samples a larger conformational space, thereby destabilizing the receptor conformation (**Figure 4D-F**). This observation is in agreement with Sakae *et al.,* who reported that the presence of fucose residues induces conformational rearrangements of both the protein environment and the N162 glycan, resulting in increased glycan mobility^29^. Our findings suggest that the detrimental effects of core fucosylation arise not solely from local steric interference at the glycan interface, but from indirect conformational and dynamical perturbations that propagate throughout the Fc–CD16a assembly.

Our analyses reveal that the impact of fucose extends beyond simple steric interference between glycans and instead propagates through larger-scale structural and dynamical changes within the Fc–CD16a assembly. Protein–Protein contact analyses revealed that dual fucosylation perturbs the overall packing and organization of the binding interface by reducing intermolecular contacts relative to their afucosylated counterparts (**Figure 5**). However, because contact maps alone provide only geometric information, we next examined how these structural differences manifest energetically. Residue-wise energy decomposition analyses revealed a striking redistribution of energetically important residues in the fucosylated systems. In the paired afucosylated systems, the dominant energetic contributions remain concentrated near the Fc–CD16a binding interface. In contrast, the paired biantennary fucosylated systems display energetically active residues extending farther into the distal D1 domain of CD16a (**Figure 6**), suggesting that the effects of fucose propagate beyond the immediate glycan environment and alter the global conformational organization of the receptor. This observation points toward a long-range structural coupling to the reduced binding affinities observed in the fucosylated systems. The emergence of energetically important residues within the distal D1 domain further suggests that the structural effects of fucosylation extend well beyond the immediate binding interface.

To further investigate these conformational effects, we constructed FES landscapes to characterize the dominant conformational states sampled by the complex during the simulations. The paired biantennary afucosylated systems preferentially occupy compact and highly stable conformational basins, whereas the fucosylated systems sample broader and more diffuse low-stability states (**Figure 7**), indicating increased conformational heterogeneity upon fucosylation. This increased conformational diffusivity is consistent with a less optimized receptor-binding geometry and a broader ensemble of weaker binding-competent states, highlighting how glycan composition can reshape the receptor-binding energy landscape through modulation of conformational populations. Because such conformational shifts could alter coordinated motions across the binding assembly, we examined residue-level dynamic correlations using DCCM analysis, which revealed that afucosylated systems exhibit substantially stronger correlated motions both within the Fc domains and across the Fc–CD16a interface (**Figure 8**). In particular, correlations between chain N of the Fc and the CD16a D2 domain are markedly enhanced in the afucosylated systems, consistent with tighter dynamic coupling at the binding interface. Together, these findings support a model in which fucosylation disrupts long-range dynamic communication within the Fc–CD16a assembly, weakening coordinated motions that stabilize binding, while afucosylation restores these long-range coupled motions and promotes stronger receptor engagement.

While our simulations provide atomistic insight into the structural and dynamical consequences of Fc fucosylation, the calculated binding energetics remain dependent on the underlying force-field representation and MM/PBSA approximations. Consequently, the energetic trends reported here are most appropriately interpreted in a comparative and mechanistic context rather than as absolute binding free energies. Furthermore, while the simulations span the microsecond timescale, they are initiated from a single experimentally resolved Fc–CD16a structure and therefore may not exhaustively capture the full conformational landscape accessible to these highly dynamic glycoprotein assemblies. This study focused on paired fucosylated and afucosylated Fc glycoforms to capture the maximal contrast in fucose-dependent modulation of Fc–CD16a binding. Asymmetric systems, in which one arm carries a fucosylated glycan and the other carries an afucosylated glycan, were not explored. Such heterogeneous glycoforms can also be biologically relevant, and extending this framework to such partially fucosylated glycoforms is a possible future direction for determining how intermediate fucosylation states tune the Fc–CD16a conformational and energetic landscape. Despite these limitations, the convergence of evidence from contact analyses, energetic decomposition, conformational landscapes, and correlated dynamics collectively supports a robust mechanistic framework for understanding how Fc fucosylation regulates Fc–CD16a binding.

In conclusion, our study provides a molecular and dynamical framework for understanding how Fc fucosylation modulates IgG1 Fc–CD16a binding. While previous experimental studies have firmly established that core fucosylation reduces binding affinity, the structural origins of this effect have remained difficult to resolve due to the heterogeneous and highly dynamic nature of glycan-mediated interactions. Through extensive atomistic molecular dynamics simulations, we show that the effects of fucose can extend far beyond local glycan rearrangements and instead propagate through broader conformational and dynamical changes across the Fc–CD16a assembly. Our results describe a substantial impact of fucosylation on the binding assembly, leading to reduced intermolecular contacts, redistribution of energetically important residues away from the binding interface, and destabilization of compact receptor-bound conformations. The afucosylated systems maintain tighter intermolecular packing, stronger dynamic coupling across the Fc–CD16a interface, and more stable conformational basins that collectively favor stronger binding interactions. Importantly, these observations suggest that the influence of fucose is not limited to direct steric effects at the glycan interface but instead alters the global conformational organization and coordinated motions of the receptor–antibody complex. More broadly, this work highlights how subtle changes in Fc glycosylation can reshape the energetic and dynamical landscape of antibody–receptor recognition. These findings further emphasize the importance of controlling Fc glycan heterogeneity during therapeutic antibody production, where even subtle variations in glycoform populations may significantly alter Fc receptor engagement and downstream immune effector responses. By resolving these effects at atomistic resolution across multiple glycan architectures, our study provides mechanistic insights into how glycan composition tunes Fc-mediated immune recognition and offers a structural framework for the rational glycoengineering of next-generation antibody therapeutics with enhanced effector functions. More broadly, our results demonstrate how glycan modifications can be exploited to rationally engineer biomolecular recognition landscapes by tuning conformational ensembles, inter-domain communication, and dynamic stability in therapeutic glycoproteins.

## METHODS

### Model setup

The crystal structure of the IgG1 Fc–CD16a complex (code: 1E4K) was obtained from RCSB^19^. The Fc domain of the complex is glycosylated for both cases of fucosylation and afucosylation with increasing degrees of galactosylation (G0, G1 and G2) (**Figure 2A,C,E**). The glycans on the CD16a were modeled based on the most probable glycans obtained from the mass spectroscopy data from Kashyap et al., with the N162 glycan replaced with High mannose (M5) to elicit the strongest binding^9, 12^ (**Figure 2C,E**). The final list of all the systems is shown in **SI Table 1**. These glycans were added to the crystal structure using ALLOSMOD, an ab initio modelling tool^35^, with the CHARMM36m forcefield^36^. Our final structure consists of chain N and chain F of the Fc domain of the IgG1, with 6 different glycosylation types bound to the D1 and D2 domains of the CD16a, with the most probable glycans modelled at positions N38, N45, N74, and N169. The N162 glycan is represented by High mannose (M5) to elicit the strongest binding^9, 12^ All visualizations and structural representations were done using VMD 1.9.3^37^ and PyMOL^38^.

### Simulation protocol

We had a total of 6 distinct systems as represented in **SI Table 1**. A transferable intermolecular potential with 3 points (TIP3P) water model was used for simulations, and a cubic water box was used as the simulation box with a padding of 20 Å to account for the motion and possible extension of glycans. The systems were neutralized at a physiological ionic concentration of 150 mM of KCl. For all these systems, minimization was performed using NAMD with a gradual release of constraints on the glycan-heavy atoms and protein backbone using a combination of conjugate gradient and steepest descent. The first equilibration step involved a restrained simulation in a constant volume and temperature (NVT) ensemble for 2 ns. The second step involved a restrained simulation with constant pressure and temperature (NPT) for 10 ns. During the first and second stages, harmonic position restraints were imposed on all non-hydrogen atoms. The third step of equilibration involved 10 ns of simulation in the NPT ensemble and only included position restraints on protein backbone atoms.

After this equilibration protocol, we ran three separate trajectories for each system, each around 1.2 μs in length. Temperature was maintained at 310 K using a Langevin thermostat (canonical ensemble sampling). Pressure was maintained at 1 atm using a Berendsen barostat^39^. This combination was used for all production trajectories. Upon examining the RMSD plots, the first 200 ns of each simulation were discarded to account for the equilibration of the system (see **SI Figure 1**). Hence, the total runtime used for all the analysis was 3 μs for each system. Since we had access to multiple graphical processing units (GPUs) on the high-performance computing (HPC) cluster, we converted our input files to a format compatible with the AMBER simulation package^24^, which provides the best throughput on GPUs for our simulations. We also used the hydrogen mass repartitioning feature in AMBER, which helps to speed up the simulations by using a timestep of 4 fs

### Binding energy calculation

The binding energy calculations were done using the MMPBSA.py package^23^ of AMBER^24^. The frames corresponding to the last 600 ns of each trajectory were selected to account for stability for a total of 1800 frames across three trajectories. We estimated relative binding affinities using an empirically corrected MM/PBSA-type approach, in which the van der Waals and electrostatic contributions are retained but scaled by literature-derived coefficients (α = 0.158 for vdw and β = 0.153 for electrostatic, respectively) to reduce the well-known tendency of single-trajectory continuum-solvent methods to overestimate absolute interaction energies, especially for buried charged contacts and protein–protein interfaces, following the approach of Minghui et al ^25^. This strategy preserves the computational efficiency and comparative interpretability of MM/PBSA while bringing the resulting energies onto a more experimentally realistic scale, making it particularly suitable for relative comparisons across different glycoforms on the same protein pairs, instead of trying to calculate exact absolute binding free energies. The use of small scaling factors for these terms is also conceptually consistent with earlier linear interaction energy (LIE)-like formulations, where raw interaction energies are empirically damped to account for solvent reorganization and structural relaxation not explicitly captured in simplified free-energy models^40^. The average values of the Van der Waals and Electrostatic energy components were then taken across these systems, along with their standard errors.

### Analysis of glycan-glycan interactions

Interactions between the two N297 glycans on the Fc were quantified by measuring contact distances between heavy atoms of the two glycans. A contact was defined when the distance between any pair of heavy atoms from the two glycans fell below a predefined cutoff of 6.0 Å. This cutoff selection is consistent with standard heavy-atom contact definitions used in structural analyses^41^ and prior Fc glycan simulation studies^42^. The minimum heavy atom distance between glycan chains was computed per frame using all pairwise heavy atom combinations, and the average minimum distance across the trajectory was used to assess differences in glycan–glycan proximity between paired biantennary fucosylated and afucosylated systems. Contact frequency values within 6.0 Å were considered favorable contacts. To investigate potential indirect effects caused by fucose on glycan and protein conformations, and given prior evidence that the N162 glycan on the CD16a plays a key role in binding, we further characterized the conformational dynamics of the CD16a N162 glycan using free-energy landscape analysis. Free Energy Surface (FES) landscapes were computed using two reaction coordinates: (i) the radius of gyration (Rg) of the N162 glycan, and (ii) the distance between the centre of mass (COM) of the N162 glycan and the COM of the Fc N297 glycans, describing its position relative to the Fc. Probability distributions along these coordinates were extracted from the MD trajectories and converted to FES plots.

### Analysis of protein-protein contacts

To analyze the contacts made between the Fc and the CD16a proteins, we measured the contacts made between the two systems. This was performed using the g_contacts module of GROMACS, using a default cutoff of 3Å^43, 44^. To better understand the differences between the Paired biantennary afucosylated and fucosylated systems, the contact frequencies under 0.25 were discarded. Additionally, the differences between the afucosylated and fucosylated frequencies were calculated and plotted to better represent the differences between the two systems.

### Residue-wise energy decomposition analysis

Per-residue energy decomposition analysis was performed to identify residues contributing most significantly to the Fc–CD16a binding interface in both fucosylated and afucosylated systems. The decomposition analysis was performed using the MMPBSA.py package^23^ of AMBER^24^. A portion of the trajectories was taken as the input, specifically the frames corresponding to the time between 600 and 1200 ns, yielding a total of 50 evenly spaced frames for each system. Binding free energy contributions were decomposed on a per-residue basis, and the top 10 most energetically significant residues at the interface were identified for each condition. To enable direct comparison between paired biantennary fucosylated and afucosylated complexes, residues that were uniquely enriched in each system were selected and explicitly represented on the Fc–CD16a structures. This mapping was used to visualize and quantify differences in the spatial distribution of energetically critical residues across the interface.

### Analysis of the motion of the CD16a relative to the Fc

To characterize the impact of fucosylation on the relative motion of CD16a with respect to the Fc domain, we quantified both rotational and translational degrees of freedom of CD16a and the Fc. This was calculated by measuring the relative orientation of CD16a and was described using two principal axes of rotation (**Figure 7G-H**). In addition to rotational motion, translational changes (or separation) within the two Fc arms were monitored by measuring the distance between the Centre of mass (COM) of the two CH2 domains. These reaction coordinates were chosen to capture coupled motions between CD16a reorientation and Fc conformational changes.

The conformational landscapes associated with these motions were quantified using two-dimensional Free Energy Surface (FES) plots. Probability distributions along each pair of reaction coordinates were extracted from equilibrated simulation trajectories and converted to free-energy surfaces. In these FES representations, deeper energy basins correspond to more stable and frequently sampled conformational states.

### Analysis of Fc–CD16a correlated motions

Dynamic cross-correlation (DCC) analysis was used to probe the correlative motions of atoms in the Fc and the CD16a. This correlative motion between two atoms *i* and *j* is defined as

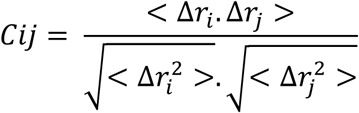

where *r_i_*(*t*) denotes the vector of the *i*th atom’s coordinates as a function of time *t*, < >_t_ is the time ensemble average and Δ*r_i_*(t) = *r_i_*(*t*) − < *r_i_*(*t*)>*_t_*. The difference in the values for the afucosylated and fucosylated systems is then plotted in the form of an *N* × *N* heatmap, where *N* is the number of Cα atoms in the system. The correlation values are calculated between −1 and 1, where 1 = complete correlation; −1 = complete anti-correlation; 0 = no correlation. The DCC matrix was generated using the MD-Task^33^ Python library and plotted using Matplotlib^45^.

## Supplementary material

The Supplementary material contains figures that are relevant to the findings in the Main manuscript. These include the complete list of systems and the RMSD plots. Filename: “**Supplementary Information**”

## Funding and Acknowledgements

N.M. and S.C. were supported by NIH NIGMS grant R35GM151231-01 and through NEU Faculty Startup Funds. N. M. acknowledges the LEADERS fellowship program jointly by Northeastern University and Amgen. This research used computational resources from Northeastern Discovery cluster at MGHPCC. The authors also acknowledge the initial tests performed by Raghavendran Suresh, which led to the start of this project.

## Author contributions

N.M. A.P. and S.C. conceptualized the study and designed the experiments. N.M. and S.C. prepared the initial structures. N.M. ran the simulations. N.M. performed the data analysis. N.M., A.P. and S.C. performed data interpretation. N.M. prepared the figures. N.M, A.P. and S.C wrote and edited the manuscript. S.C. is the corresponding author.

## Data and materials availability

Simulation trajectories and data are shared for open access in Mendeley Data (V1, doi: 10.17632/zxzmfdmz4y.1). Further materials and simulation codes are available from the corresponding authors upon request.

