## Supplemental Information Tables and Figures for "Molecular Basis of Core Fucosylation-Dependent Modulation of IgG1 Fc–CD16a Binding"

### Affiliations

| System ID | Glycan on chain N | Glycan on chain F | Type |
|---|---|---|---|
| a2g0-a2g0 | a2g0 | a2g0 | Paired |
| a2g0f-a2g0f | a2g0f | a2g0f | Paired |
| a2g1-a2g1 | a2g1 | a2g1 | Paired |
| a2g1f-a2g1f | a2g1f | a2g1f | Paired |
| a2g2-a2g2 | a2g2 | a2g2 | Paired |
| a2g2f-a2g2f | a2g2f | a2g2f | Paired |

**SI Figure 1**: Overview of glycan systems. The table shows the 6 different glycan structural variants analyzed in our study. Chain N and Chain F represent the CD16a proximal and distal arms on the Fc, respectively.

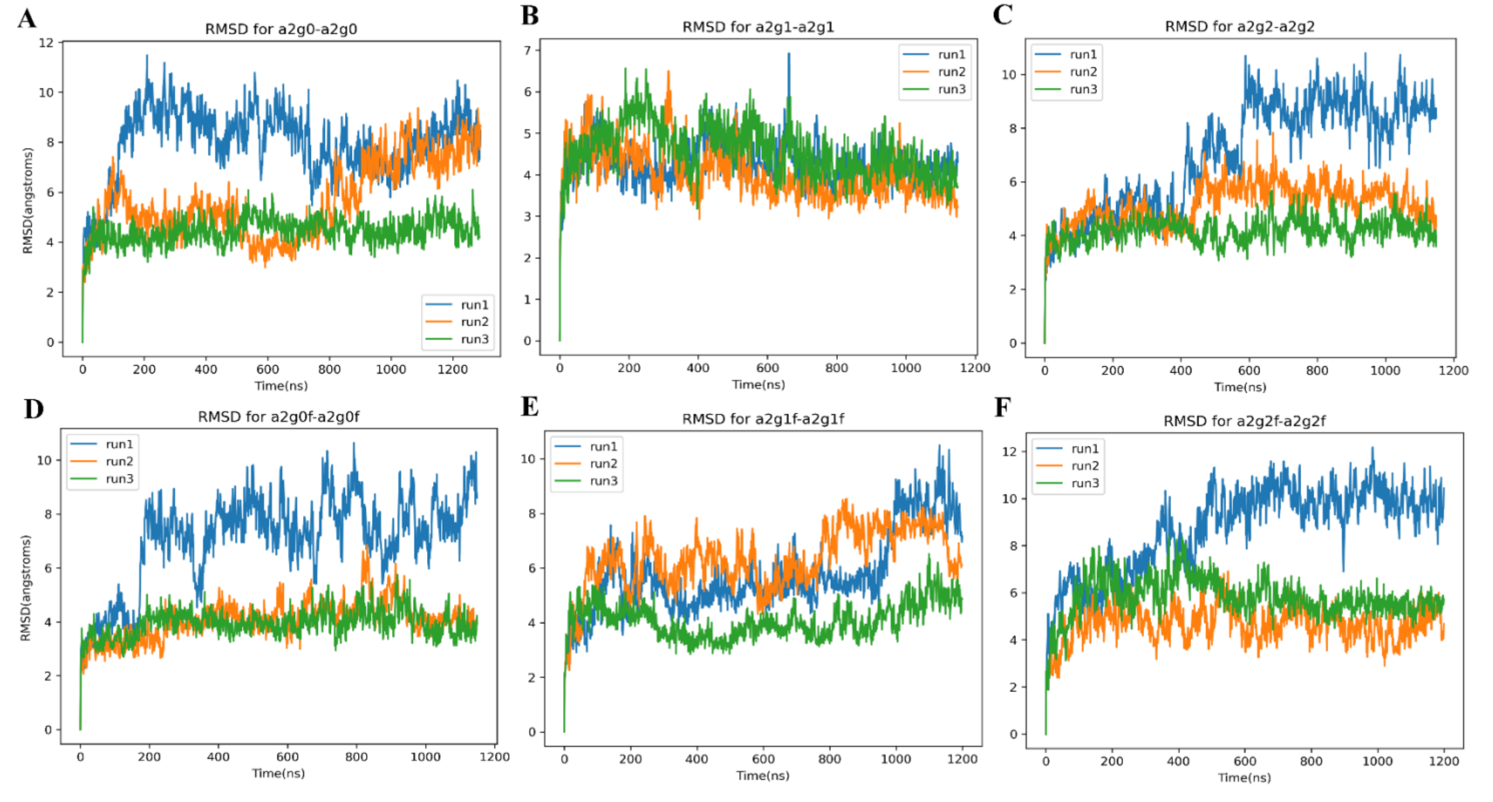


**SI Figure 2**: RMSD plots for the dual afucosylated (**Panels A, B** and **C**) and dual fucosylated (**Panels D, E** and **F**), respectively. Each of run1, run2 and run3 corresponds to individual trajectories for each system.

| System | Residue on the Fc | Location on the Fc | Residue on the CD16a | Location on the CD16a |
|---|---|---|---|---|
| a2g0-a2g0 | LYS94 | Chain F | PHE130 | D2 |
| a2g0-a2g0 | ILE104 | Chain F | ANS112 | D2 |
| a2g0-a2g0 | PRO4 | Chain F | LEU115 | D2 |
| a2g0-a2g0 | PRO226 | Chain N | LEU17 | D1 |
| a2g0-a2g0 | LYS310 | Chain N | ASN133 | D2 |
| a2g0f-a2g0f | PRO10 | Chain F | GLU65 | D1 |
| a2g0f-a2g0f | GLU221 | Chain N | ARG152 | D2 |
| a2g0f-a2g0f | ALA315 | Chain N | GLU82 | D1 |
| a2g0f-a2g0f | GLN44 | Chain F | TYR129 | D2 |
| a2g0f-a2g0f | LEU223 | Chain N | LYS158 | D2 |
| a2g1-a2g1 | PRO220 | Chain N | GLU18 | D1 |
| a2g1-a2g1 | GLY9 | Chain F | VAL160 | D2 |
| a2g1-a2g1 | ASN285 | Chain N | ASN133 | D2 |
| a2g1-a2g1 | THR287 | Chain N | PHE130 | D2 |
| a2g1-a2g1 | PRO10 | Chain F | VAL118 | D2 |
| a2g1f-a2g1f | LYS89 | Chain F | GLU43 | D1 |
| a2g1f-a2g1f | LYS62 | Chain F | ARG152 | D2 |
| a2g1f-a2g1f | PRO103 | Chain F | LYS117 | D2 |
| a2g1f-a2g1f | ASN48 | Chain F | ASP62 | D1 |
| a2g1f-a2g1f | ARG64 | Chain F | HSE131 | D2 |
| a2g2-a2g2 | ASP253 | Chain N | ASN133 | D2 |
| a2g2-a2g2 | ASP364 | Chain N | ALA58 | D1 |
| a2g2-a2g2 | ASP258 | Chain N | GLN122 | D2 |
| a2g2-a2g2 | ASP387 | Chain N | LEU115 | D2 |

| | | | | |
|---|---|---|---|---|
| a2g2-a2g2 | GLU221 | Chain N | SER47 | D1 |
| a2g2f-a2g2f | GLU105 | Chain F | HSE131 | D2 |
| a2g2f-a2g2f | GLU202 | Chain F | LYS25 | D1 |
| a2g2f-a2g2f | GLU129 | Chain F | GLY156 | D2 |
| a2g2f-a2g2f | GLU154 | Chain F | LYS128 | D2 |
| a2g2f-a2g2f | LYS314 | Chain N | LYS158 | D2 |

**SI Figure 3:** The top five energetically active residues identified at the Fc–CD16a interface for each glycoform complex, categorised by Fc chains and CD16a domains. The entire list can be provided upon request
